# Dynamic fMRI networks of emotion

**DOI:** 10.1101/2025.02.05.636581

**Authors:** Niels Janssen, Uriel K.A. Elvira, Joost Janssen, Theo G.M. van Erp

## Abstract

The experience of emotions is that of dynamic, time changing processes. Yet, many fMRI studies of emotion average across time to focus on maps of static activations, overlooking the temporal dimension of emotional responses. In this study, we used time-resolved fMRI, group spatial independent component analysis (ICA), dual regression, and Gaussian curve fitting to examine both the spatial and temporal properties of whole-brain networks during a behavioral task. This task included trials that spanned over 25 seconds of watching short, emotionally evocative movie clips, making emotion-related decisions, and an intertrial rest period. We identified four whole-brain networks with unique spatial and temporal features that mapped onto different stages of the task. A network activated early in the course of the task included perceptual and affective evaluation regions, while two later networks supported semantic interpretation and decision-making, and a final network aligned with default mode activity. Both spatial and temporal properties of all four networks were modulated by the emotional content of the movie clips. Our findings extend current models of emotion by integrating temporal dynamics with large-scale network activity, offering a richer framework for understanding how emotions unfold across distributed circuits. Such temporal-spatial markers of emotional processing may prove valuable for identifying and tracking alterations in clinical populations.

## Introduction

The relationship between emotions and brain function has been a central focus of affective neuroscience, with foundational studies elucidating the roles of structures such as the amygdala, insula, and prefrontal cortex in emotional processing (LaBar, LeDoux, Spencer, & Phelps, 1995; Ochsner, Bunge, Gross, & Gabrieli, 2002). Despite significant progress, most functional MRI (fMRI) studies have adopted a static view of emotions, often emphasizing localized brain activations or static connectivity patterns rather than the dynamic and distributed nature of emotional processing (see Kuppens & Verduyn, 2017; Pessoa, 2017; Waugh, Shing, & Avery, 2015, for discussion). This particular approach to studying emotions arises in part because analytical techniques collapse time-varying fMRI activity into static spatial maps and also because experimental paradigms typically use unchanging stimuli (Murphy, Nimmo-Smith, & Lawrence, 2003; Phan, Wager, Taylor, & Liberzon, 2002). In the present study, we collected fMRI data while participants performed a behavioral task with trials that paired the viewing of emotionally evocative short movie clips with a subsequent discrete emotion decision task as well as an intertrial rest period. We sought to address three key questions: (1) What distinct whole-brain networks underlie performance during this task? (2) Do the spatial distributions of these networks vary as a function of the emotional content of the stimuli? (3) Are the temporal dynamics of these networks—such as their time to peak, peak value or duration sensitive to the emotional characteristics of the stimuli? To answer these questions we relied on a data-driven approach that emphasized a combined spatial and temporal view on fMRI data analysis.

Previous studies investigating emotional processing with fMRI have generally revealed a static view of emotions in the brain. Specifically, recent research has highlighted that emotions engage widespread networks involving multiple interconnected regions, reflecting the complex and distributed nature of emotional processing (Kober et al., 2008; K. A. Lindquist, Wager, Kober, Bliss-Moreau, & Barrett, 2012; Palomero-Gallagher & Amunts, 2022; Pessoa, 2017; Zhou et al., 2021). For instance, Riedel et al. (2018) applied a meta-analytic clustering approach to over 1,700 neuroimaging experiments to delineate five distinct whole-brain networks associated with emotional processing. The first two networks were linked with visual and auditory perception, with convergent activation in the visual (occipital) and auditory (superior temporal) cortices, respectively. The third network captured the salience network, including the insula and dorsal anterior cingulate cortex, reflecting attention to emotionally salient information. The fourth network was associated with appraisal and prediction of emotional events, involving regions of the default mode network, such as the medial prefrontal and posterior cingulate cortices. The fifth network focused on the induction of emotional responses, with key contributions from the amygdala, parahippocampal gyri, and fusiform gyri, underscoring its role in generating contextually relevant emotional responses. While this and other studies highlight the involvement of large scale networks in the processing of emotion and therefore go beyond the traditional one-to-one mapping between brain regions and emotions (e.g., Murphy et al., 2003), they do not reveal the temporal component associated with these networks.

Traditionally, many fMRI studies of emotion have employed decontextualized static images of facial expressions or evocative pictures to elicit emotional responses (Adolphs, 2002; Liu, Liu, Zheng, Zhao, & Fu, 2021; Wallenwein, Schmidt, Hass, & Mier, 2024). In contrast, more recent studies have argued that movies offer a richer perceptual experience that is of a higher ecological validity (Finn & Bandettini, 2021; Sonkusare, Breakspear, & Guo, 2019). Filmmakers are adept at eliciting specific emotions in viewers (Westermann, Spies, Stahl, & Hesse, 1996), and recent studies have shown that using movies as emotion-evoking stimuli can provide new insights into the relationship between emotions and the brain (Jääskeläinen, Sams, Glerean, & Ahveninen, 2021; Morgenroth et al., 2023; Saarimäki, 2021). However, a limitation of current studies using movie clips is that they often employ lengthy segments lasting several minutes, during which multiple cognitive and emotional events can occur (e.g., Vemuri & Surampudi, 2015; Xu et al., 2023). This complexity makes it challenging to disentangle the dynamic changes in emotional processing over time (but see, Meer, Breakspear, Chang, Sonkusare, & Cocchi, 2020). To address this issue, shorter movie clips focusing on a single event can be utilized to better control and observe how emotions develop over time within events. In addition, an important aspect of emotions concerns not only their initial expression but also how they are regulated following that initial response (Etkin, Büchel, & Gross, 2015; Ochsner & Gross, 2008). This suggests that the study of the brain dynamics of emotions should focus not only on the stimulus event but should also examine how brain responses change after the event has concluded. Incorporating short periods of rest after the presentation of short movie clips may facilitate the investigation of post-event emotional regulation processes and how they are represented in the brain.

In the current study, participants viewed 12.5s audiovisual clips selected from the movie Forrest Gump (Zemeckis et al., 1994). Each clip was specifically chosen to evoke a single emotion—happiness, fear, or sadness—with each emotion equally represented across a total of 15 clips. Following each clip, participants completed a two-alternative forced-choice decision task using discrete emotion words denoting the emotion experienced by the movie’s protagonist. Finally, there was a 10s intertrial rest period. We collected whole-brain fMRI data at 2mm spatial resolution, and extracted the event-related fMRI signal at 2s temporal resolution across a 28s epoch for each emotion type and for each participant using slice-based fMRI (Janssen, Hernández-Cabrera, & Foronda, 2018; Janssen & Mendieta, 2020). In parallel with resting-state analyses, we then performed group spatial Independent Component Analysis (ICA) to identify large-scale networks with distinct spatial distributions and temporal profiles (Beckmann & Smith, 2004; Smith et al., 2009). The spatial and temporal properties of the discovered networks were further quantified using dual regression (Beckmann, Mackay, Filippini, Smith, et al., 2009) and Gaussian curve fitting (Janssen & Mendieta, 2020; Kruggel, Zysset, & von Cramon, 2000; M. A. Lindquist, Loh, Atlas, & Wager, 2009), respectively. Finally, we performed mixed-effects regression modeling and inferential statistics to establish the reliability of our results.

The emotions that are experienced by the protagonist in each clip developed gradually over the course of each clip rather than appearing immediately. In other words, while the visual and auditory perception of the movie clip is instantaneous, the comprehension of its emotional content requires more nuanced processing that lags behind the immediate perceptual input stream. Therefore, we expected to detect brain networks associated with initial sensory processing, involving primary visual and auditory cortices (Riedel et al., 2018), as well as networks related to higher-level cognitive and emotional processing, such as those involving the middle temporal gyrus and other association areas (Huth, De Heer, Griffiths, Theunissen, & Gallant, 2016; Visser, Jefferies, Embleton, & Lambon Ralph, 2012). Given that we analyzed the entire behavioral task—including the decision stage and the rest period—we anticipated identifying additional large-scale networks associated with these later stages of the task, such as a response initiation network involving primary motor cortices and the default mode network (Satpute & Lindquist, 2019).

## Methods

### Participants

Twenty-five native speakers of Spanish took part in the experiment (13 females, mean age 21.9 yrs, sd 3.9 yrs). Participants were recruited from the student population at the University of La Laguna, and received course credit or were paid 10 Euros. All participants were right-handed and had no neurological or psychiatric disorders and were not taking any medication. The study was conducted in compliance with the declaration of Helsinki, and all participants provided informed consent in accordance with the protocol established by the Ethics Commission for Research of the University of La Laguna (Comite de Etica de la Investigacion y Bienestar Animal). Participants answered a brief questionnaire about their knowledge of the movie Forrest Gump. Due to technical problems and on the basis of quality control procedures for both structural and functional data (see Figures 1A and B), data from three participants were excluded from further analyses.

**Figure 1.**
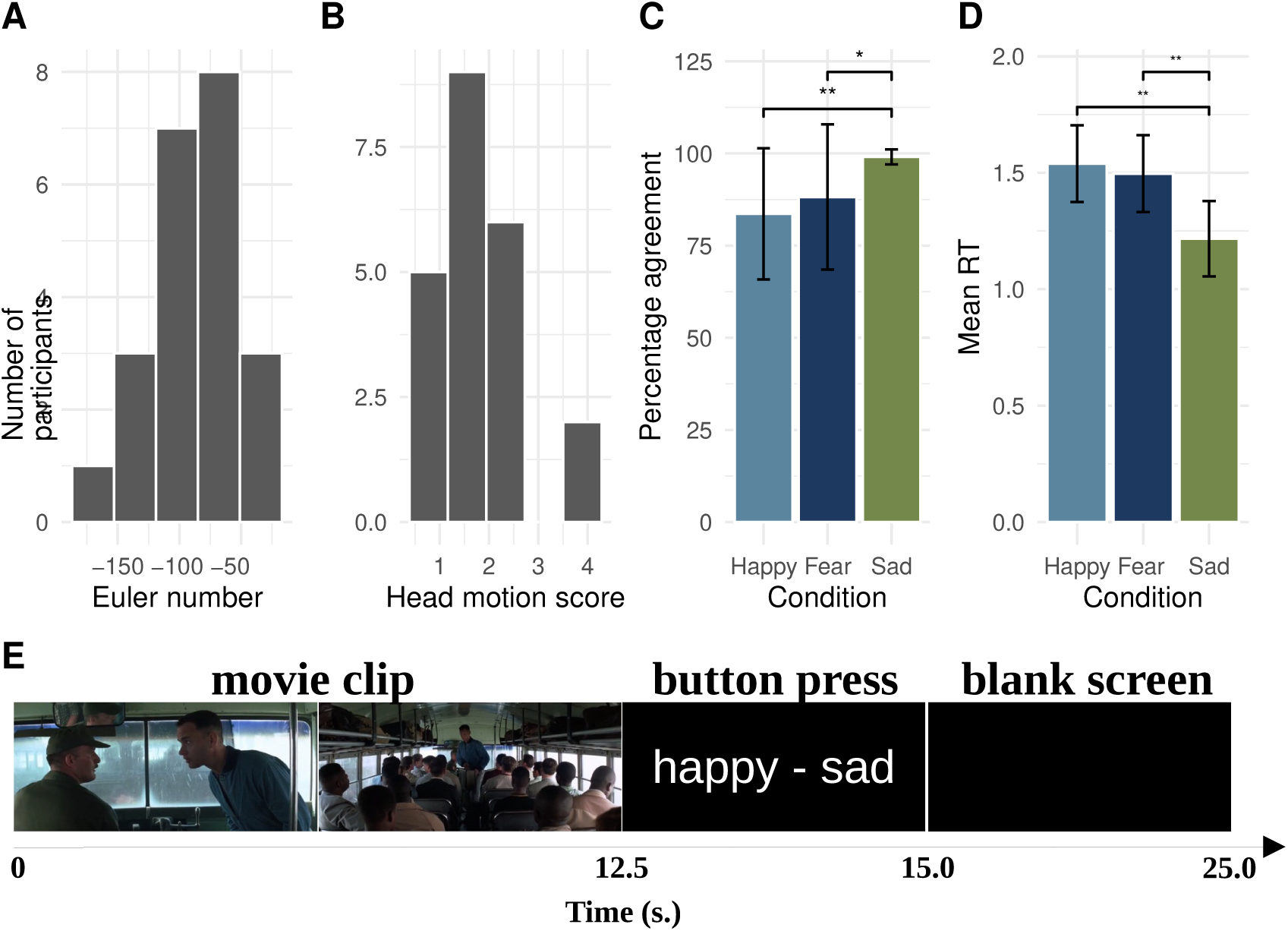
Overview of Quality Control of structural data with Euler number (A), fMRI head motion scores (B), analysis of behavioral C) and reaction times (D) and overview of the task (E).

#### Behavioral task

We selected 15 clips from the movie Forrest Gump (Zemeckis et al., 1994). The movie was dubbed with Spanish (mainland Spain) audio. Each clip was exactly 12.5 s long and was chosen such that it portrayed a happy, sad, or fearful experience for the Forrest Gump character (see Figure 1E). Clips were chosen such that they contained a natural onset and offset with no abrupt cuts and were played at their intended speed. For example, in one of the clips, Forrest walks in a garden and sees Jenny again after a long time, constituting a happy experience. The order of the fifteen video clips followed the natural story progression of the movie and therefore did not vary between participants. Please see Table 1 for the specific order of the emotions expressed by Forrest across trials. Clips were presented with sound. To ensure the sound was audible over the scanner noise, participants wore headphones and we used the ffmpeg tool to boost the sound of the clips by 300%. Pilot testing ensured the sound was audible.

**Table 1.** Order of the movie clips in the experiment.

| Clip# | Emotion | description |
| --- | --- | --- |
| 1 | Happy | Young Forrest meets Jenny |
| 2 | Fear | Forrest runs from bullies |
| 3 | Sad | Forrest says goodbye to Jenny |
| 4 | Fear | Forrest gets shouted at |
| 5 | Sad | Forrest is alone in bed |
| 6 | Fear | Forrest get attacked |
| 7 | Happy | Forrest sees Jenny |
| 8 | Fear | Forrest sees dangerous situation |
| 9 | Sad | Forrest sees sad person |
| 10 | Sad | Forrest sees sick Jenny |
| 11 | Fear | Forrest sees Jenny ghost |
| 12 | Happy | Forrest sees Jenny |
| 13 | Sad | Forrest knows what love is |
| 14 | Happy | Forrest is with Jenny |
| 15 | Happy | Forrest stays with Jenny |

Prior to the experiment, participants were instructed to watch each clip and then make a two-alternative forced choice on the emotion experienced by Forrest. Participants were given handheld controllers in their left and right hands and instructed to press a button with their index finger on the left or right hand following the left-right location of their preferred emotion word on the screen. On a given trial, the movie clip was presented for 12.5 s followed by two emotion words for 2.5 s, and finally followed by a 10 s blank screen (see Figure 1E for a graphical overview). The total trial time was 25 s. The total duration of the experiment was 6.25 minutes.

The experiment involved a single run but was conducted in the context of a larger acquisition protocol in which an additional resting-state scan was acquired. Acquisition of these two fMRI runs was preceded by T1w and T2w image acquisitions (see below for sequence details). Stimulus presentation was controlled by Neurobs Presentations (v14). Participants in the scanner viewed and listened to the stimuli with MRI compatible goggles and headphones made by VisuaStim. The goggles provided an image resolution of 800 by 600 pixels at 60 Hz. The MRI compatible handheld controllers were also made by VisuaStim. Stimulus presentation was directly synchronized with the MRI machine.

#### MRI acquisition

All images were acquired using a 3T Signa Excite scanner (General Electric, Milwaukee, WI, USA) with a standard transmit/receive 8 channel head coil. Head movement was minimized by fixating MRI compatible goggles between a participant’s head and the coil using spongepads. fMRI data were acquired using Gradient-Echo EPI. The sequence parameters were chosen to provide high spatial-resolution while preserving whole brain coverage. We used an axial zoomed-acquisition where our FOV and acquisition matrix were reduced in the (left-right) phase encoding direction (Olman, Davachi, & Inati, 2009). Specifically, the slice thickness was 2.6 mm (no gap), FOV was 256 x 128 mm and the matrix was 128 x 64 resulting in 2 x 2 x 2.6 mm voxels. To achieve whole-brain coverage we collected 39 slices and set the TR to a relatively long 3 s. However, because stimulus onset was jittered with respect to the TR, as explained below, the value of the TR does not determine the temporal resolution of the extracted fMRI signal (Janssen et al., 2018; Josephs, Turner, & Friston, 1997). The echo-time was 33 ms and the flip angle 90 degrees. In the fMRI run 130 volumes were collected and lasted 6.5 minutes.

Finally, to comply with Human Connectome Project minimal preprocessing pipeline requirements (Glasser et al., 2013), both T1w and T2w whole-brain structural images were acquired. T1w images were obtained using the 3D FSPGR with ASSET sequence: TI/TR/TE: 650/8.8/1.8 ms, flip angle = 10 degrees, 196 slices, FOV 256 x 256 mm, slice thickness 1 mm, matrix 256 x 256 resulting in 1 mm isotropic voxel sizes. T2w images were acquired using a spin-echo sequence: TR/TE: 15000/106 ms, NEX 2, prescribing 110 sagittal slices, FOV 256 x 256 mm, slice thickness 1.5 mm, matrix 256 × 512 (with 2x parallel imaging acceleration) resulting in 0.5 x 0.5 x 1.5 mm voxels.

#### Preprocessing

Preprocessing of structural and functional data relied on the Human Connectome Project (HCP) minimal preprocessing pipeline (v4.7.0; Glasser et al., 2013). For the preprocessing of the T1 and T2w images we used the pre-freesurfer and freesurfer batch scripts with options that were applicable to our data (i.e., no readout distortion correction, 1 mm MNI HCP structural templates, no fieldmap correction). Freesurfer v6.0 was installed on our system (Fischl, 2012). This step therefore produced cleaned T1 and T2w images that were then fully processed using the Freesurfer v6.0 program. From the freesurfer output we were particularly interested in the volumetric cortical and subcortical segmentation produced in the aparc+aseg file. Note this file was produced in native T1w space by the pipeline.

For the preprocessing of the fMRI data we used the generic fMRI volume batch script from the HCP preprocessing pipeline. All registrations (EPI to T1w and T1w to MNI) were manually checked for inconsistencies (none were detected). Next, the functional data were temporally filtered at 2000 s and manually cleaned using Independent Component Analysis (Griffanti et al., 2014). We obtained Independent Components for all fMRI datasets using FSL Melodic v3.15 with default settings (i.e., automatic estimation of the number of components; Beckmann & Smith, 2004). An experienced rater of Independent Components (NJ) then hand-classified all components as signal or noise for all participants using in-house tools (https://github.com/iamnielsjanssen/display_melodics). Components identified as noise were then regressed out of the data using the FSL regfilt tool. In the end we obtained the cleaned fMRI datasets for each participant in native fMRI space that were bias corrected, motion corrected, brain-extracted, intensity corrected, temporally filtered and ICA-cleaned. Quality control of the structural data was performed by computing Euler’s number from the freesurfer output (Rosen et al., 2018), and by computing for all functional data a composite score of head motion displacements from the six motion regressors (Jenkinson, Bannister, Brady, & Smith, 2002, see Figure 1A and B for a graphical presentation).

### Analyses

#### fMRI signal extraction

The first step in the analyses consisted of extracting the event-related fMRI signal using slice-based fMRI (see Figure 2A and B for a graphical explanation; Janssen et al., 2018). The slice-based fMRI technique offers a significant advancement over traditional fMRI analysis methods by providing improved temporal precision and flexibility in extracting the fMRI signal. FMRI data is typically acquired by sequentially sampling different sections of the brain (slices) over a single repetition time (TR). In standard fMRI analyses, whole-brain volumes are constructed from all slices acquired within a single TR, which is well-known to introduce temporal distortions (Parker, Liu, & Razlighi, 2017; Sladky et al., 2011). In contrast, the slice-based technique avoids these temporal distortions by creating whole-brain volumes that are composed out of slices that are acquired at the same moment in time *relative to a presented stimulus*. A further advantage of this method is that because slices are acquired relatively rapidly, under optimal conditions, whole-brain volumes can be constructed with a temporal resolution that is equal to the TR divided by the number of prescribed slices (typically on the order of tens of milliseconds). This temporal resolution can then be flexibly adjusted by “binning” across time-points, which effectively reduces the temporal resolution while enhancing the signal-to-noise ratio (SNR). The ability to flexibly adjust the temporal resolution makes the slice-based technique especially valuable for studying the dynamics of brain activity underlying complex behavior.

**Figure 2.**
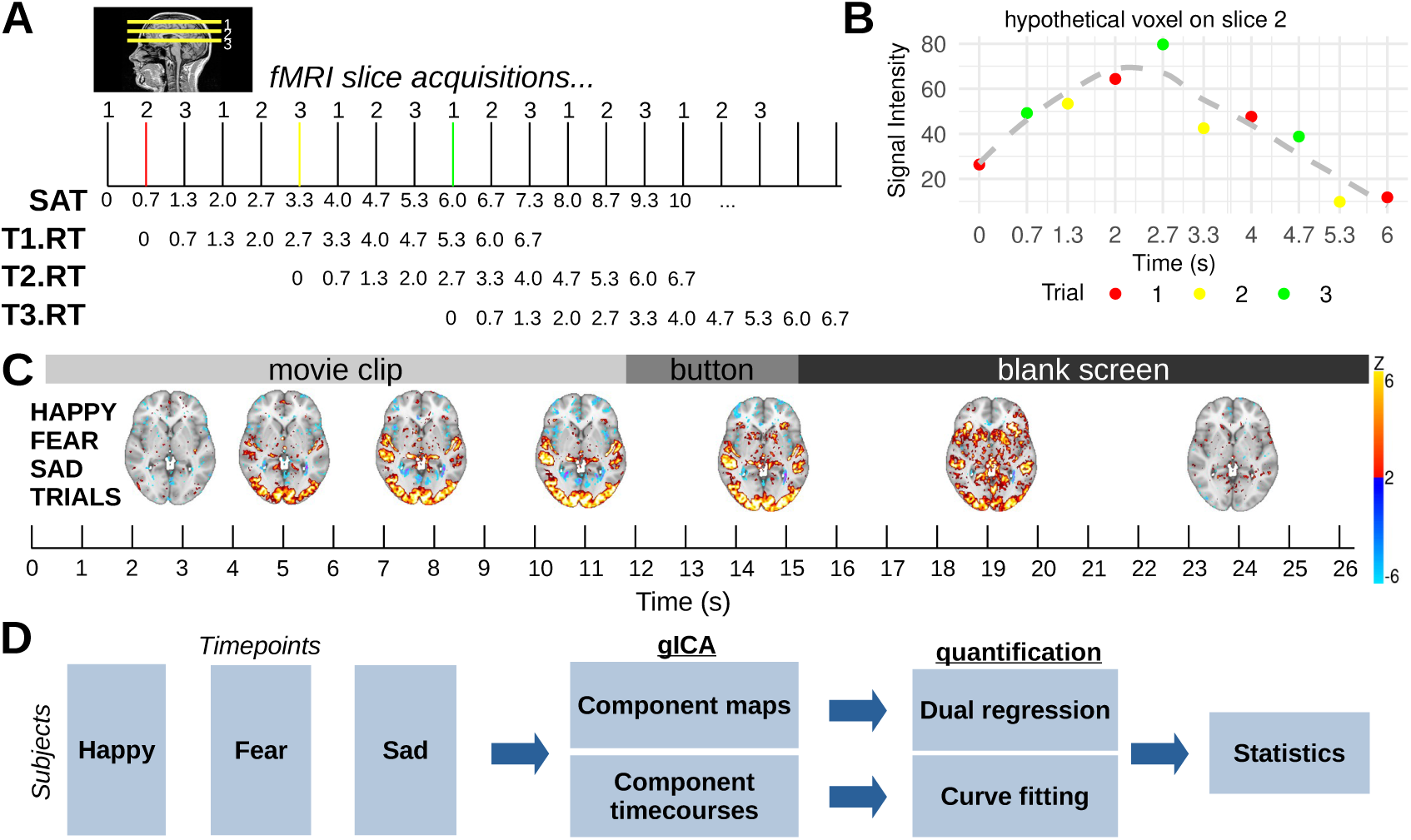
Slice-based fMRI explanation (A,B), results (C) and subsequent processing steps (D). In a hypothetical fMRI data acquisition protocol (A), three slices are acquired with TR=2s resulting in Slice-Acquisition Times (SAT) of 0, 0.7, 1.3s etc. The key point is that stimulus presentations are combined to extract the fMRI signal at high temporal resolution (Janssen et al., 2018). Three stimulus trials (T1.RT, T2.RT, T3.RT and red, yellow, green bars) whose onset coincides with each slice results in slices sampling the brain at all timepoints during a 6s epoch *relative* to the onset of the stimulus. Thus, for a specific voxel on slice 2 (B), after these three stimulus trials, signal intensities at a temporal resolution equal to the ΔSAT (0.7s) are acquired. Statistical models can extract the fMRI signal (B, grey dashed line) at this or at a higher powered binned, but lower temporal resolution. Results from group-level Slice-Based analysis of movie clips at 2s resolution for the separate happy, fear and sad trials (not all timepoints shown). Note results reveal complex increases (hot colors) and decreases (cool colors) of signals across the whole brain during the 26s epoch (C). The slice-based datasets for the happy, fear and sad trials for each subject were then concenated and entered into group spatial ICA for the detection of component maps and timecourses (D).

In the current study we used the slice-based technique to extract the fMRI signal during a 28 s epoch separately for the happy, sad and fear trials at 2 s temporal resolution. Note we chose an epoch duration that was slightly longer than the task to accommodate for some uncertainty in the offset of fMRI activities. This meant that for each voxel, the fMRI signal was extracted by comparing fMRI signal intensities collected during an interval of 3 seconds prior to stimulus onset to fMRI signal intensities at all subsequent timepoints in the 28 s epoch using a simple linear model with the variable Time as a factor (see Figure 2B for a graphical explanation). This therefore produced for every participant three 4D fMRI files that represented the event-related changes for the happy, sad and fear trials. Each file contained 14 volumes, one for every 2 seconds. Note all these 4D fMRI files were in native fMRI space.

#### Group spatial-ICA

The goal of this step was to discover the spatial and temporal fMRI patterns that arise during the behavioral task. To this end, the set of 66 4D slice-based fMRI files (22 participants with 3 experimental conditions each) were transformed to 2 mm MNI space using non-linear warping and entered into group spatial Independent Component Analysis (ICA; see Figure 2D for a graphical overview). Here we used FSL Melodic (v3.15) with default settings, except that we disabled data reduction with MIGP (i.e., we used the full concatenation). To determine the optimal number of dimensions, Melodic uses a Bayesian approach, which involves evaluating how well different models (with varying numbers of dimensions) explain the observed data while accounting for model complexity (Minka, 2000). For our data the estimated optimal number of dimensions was 6. As we will show below, from these 6 Independent Components (ICs; herein also referred to as IC networks or simply brain networks), 4 ICs were identified as signal whereas 2 ICs were identified as noise. To make such classifications we relied on relatively straightforward protocols explained elsewhere (Griffanti et al., 2014).

Importantly, spatial ICA not only outputs spatial information in the form of statistical maps but also produces an estimated timecourse that is associated with each spatial map. More formally, ICA decomposes the fMRI data matrix *X* (with dimensions corresponding to time points and voxels) into a product of two matrices: *X* = *W ^−^*^1^*M*, where *M* represents the spatially independent components (spatial maps) and *W ^−^*^1^ represents the associated time courses of these components (mixing matrix). The rows of *M* correspond to spatial patterns, which are maximally independent, while the columns of *W ^−^*^1^ contain the time series for each component. These time series describe the temporal dynamics of each spatial component. Together, the spatial and temporal components offer a complete representation of the independent processes that contribute to the observed fMRI data. In the case of FSL Melodic, the resulting melodic_mix file contains the matrix *W ^−^*^1^, where each column corresponds to the temporal activation pattern of an independent component. This temporal information allows us to examine how each spatial network’s activity fluctuates over time, providing critical insights into task-evoked brain activity. In the current study this temporal information is utilized in two ways: it is presented visually alongside the spatial map as the average across all subjects and experimental conditions (Duann et al., 2002; Janssen & Mendieta, 2020), and it is used analytically to perform Gaussian curve-fitting analyses (explained below).

#### Dual Regression

To establish the reliability of the activity of brain regions involved in these four IC networks, we performed back-projection of the group spatial ICA map using Dual Regression in native space. We then extracted regional activities for each IC network using the Desikan-Killany atlas for each individual participant (Desikan et al., 2006, see Figure 2D for a graphical overview). Reliability of these regional activities was estimated using linear mixed-effect regression modeling using the following model formula:

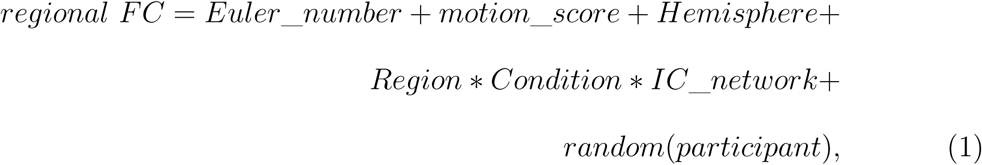

where *Euler*_*number* represented the quality control estimate for a participant’s structural data, *motion*_*score* the quality control estimate for a participant’s functional data and *Hemisphere* the functional connectivity for each hemisphere (left vs right). Although the model includes all interactions, we were particularly interested in the interactions *Region* ∗ *IC*_*network* which tested the degree to which regional functional connectivity was the same across discovered networks, and *Region* ∗ *Condition* ∗ *IC*_*network*, which examined if the aforementioned interaction depended on the emotion condition. Finally, we included a random intercept for participant to account for potential between-participant variability.

This modeling procedure therefore allowed us to establish whether a given region within a given IC network was reliably activated across participants. In other words, this would allow us to obtain a list of both cortical and subcortical brain regions that are reliably activated within each IC network. Given that the Desikan-Killany atlas is fitted by Freesurfer to the native morphology of an individual’s brain, we improved spatial precision by performing these analyses in native fMRI space.

#### Curve-fitting

The final step involved quantifying the temporal activation dynamics of the independent components (ICs). As mentioned earlier, Melodic outputs the temporal information associated with each identified IC network. Since the input to the group spatial ICA was the 4D slice-based fMRI data for each participant and experimental condition, the resulting time course information was available for each IC network, participant, and condition. These time courses were then consolidated into a single dataset with the variables Participant, IC Network, Emotion Condition, and Time, facilitating further analysis of temporal activation patterns across experimental conditions. We then performed nonlinear regression (Janssen & Mendieta, 2020; Kruggel & von Cramon, 1999; Kruggel et al., 2000) for all combination of variables using a Gaussian function of the form:

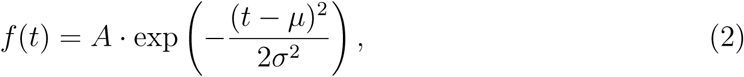

where *A* represents the amplitude, *µ* the mean and *σ* the standard deviation. As can be seen in Figure 7A, these parameters can be used to compute the peak value (*A*), the time to peak (*µ*) and duration (the FWHM equal to 2.355 ∗ *σ*) of a fitted Gaussian function. Goodness of Fit (adjusted *R*^2^) values can also be computed as a difference between observed and fitted datapoints. Gaussian fitting was performed in R using the package minpack.lm v1.2 (Elzhov, Mullen, Spiess, & Bolker, 2022), which fits a function to a set of datapoints using Levenberg-Marquardt nonlinear least squares algorithm. Initial values were estimated from the data.

These curve-fitting analyses provided amplitude, time to peak and duration estimates for each participant, Emotion Condition and IC network. Finally, statistical analyses were used to test whether peak amplitude, time to peak, and duration varied as a function of IC network and Emotion Condition using a formula of the form:

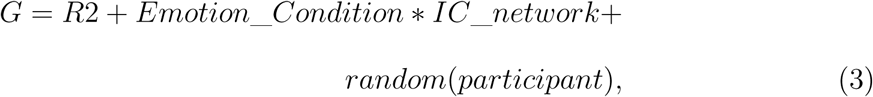

where *G* referred to the peak amplitude, time to peak, or duration. *R*2 (a Goodness of Fit value) was included in the analyses as a control variable. Here we were particularly interested in the main effect of *IC*_*network* which would reveal whether peak value, time to peak and duration values were equal for the four IC networks. In addition, we were interested in the *Emotion*_*Condition* ∗ *IC*_*network* interaction, which would reveal whether peak amplitude, time to peak or duration estimates would be equal between the three emotion conditions (happy, sad, fear) across all IC networks.

All statistical modeling reported in the current study was performed in R v4.1.2 using the packages lme4 v1.1, lmerTest v3.1, and emmeans v1.8.4 (Bates, Mächler, Bolker, & Walker, 2015; Kuznetsova, Brockhoff, Christensen, et al., 2017; Lenth, Singmann, Love, Buerkner, & Herve, 2018). Note that to reduce complexity we controlled rather than explored the effects of Hemisphere and that we modeled random participant variability using a random intercept. All post-hoc tests were performed with emmeans and corrected for multiple comparisons using the Bonferroni correction.

## Results

Analysis of the behavioral data showed that participants did not treat the three emotion categories in the same way. Specifically, as can be seen in Figure 1C, there was higher agreement among participants on the target response in the sad vs the other two emotion categories (Sad vs Happy *p <* 0.02; Sad vs Fear *p <* 0.04). In addition, as can be seen in Figure 1D, decisions for the sad condition were faster compared to the other two conditions (Sad vs Happy *p <* 0.003; Sad vs Fear *p <* 0.008).

Group spatial ICA of the 4D slice-based fMRI datasets first revealed that the estimated optimal model selection occurred with 6 dimensions. As can be seen in Figure 3, four of these networks corresponded to signal, two were considered noise. Figure 3 shows both spatial distributions and mean temporal profiles associated with each IC network. Visual inspection of these temporal profiles enables an association with the different stages of the behavioral task. Specifically, IC0 and IC1 show an activation pattern consistent with the movie watching task where there was an immediate onset, a sustained activity and offset consistent with the presentation of the 12.5 s movie clips. Similarly, IC2 shows a pattern associated with the emotion decision stage with a peak in the decision-stage of the task. In addition, the spatial distribution of co-activities of IC4 are consistent with the default mode network (dmn). For expository reasons we labeled the IC networks in terms of a tentative functional interpretation. Specifically, for reasons we discuss in more detail below, we labeled IC1 with “input”, IC0 with “meaning”, IC2 with “response” and IC4 with “dmn”. We also ordered these IC networks on the basis of their assumed temporal order of activation (i.e., IC1→IC0→IC4→IC2. Finally, IC3 and IC5 were classified as noise due to the presence of “activation” in non gray-matter areas of the brain (CSF, veins; Griffanti et al., 2014).

**Figure 3.**
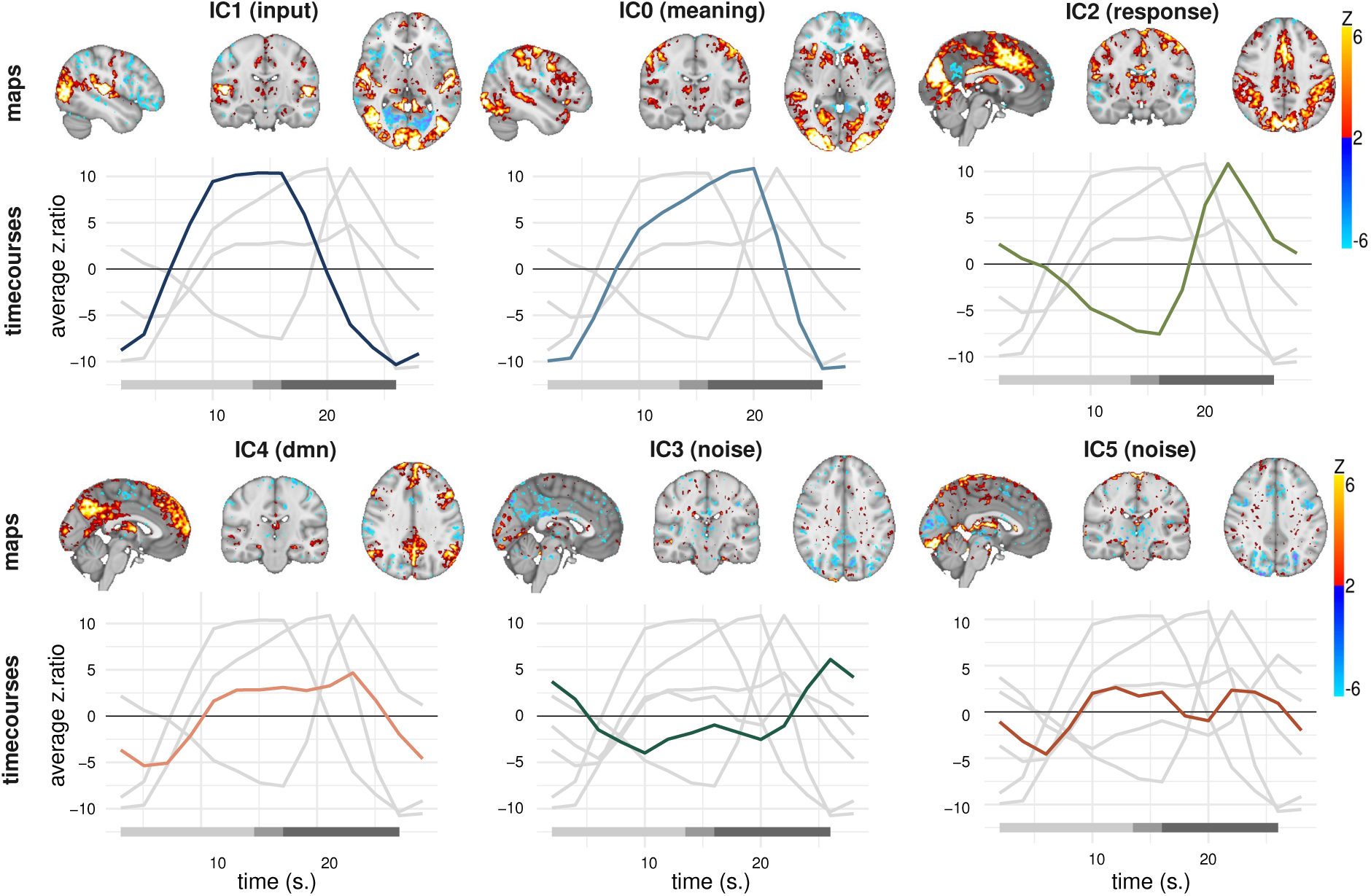
Results from group spatial ICA with 6 dimensions. Plotted are both the spatial maps as well as the component time courses averaged across subjects and conditions associated with each IC. Consideration of both spatial and temporal properties of these 6 ICs led to the identification of four ICs as signal (IC0, IC1, IC2, IC4) and two as noise (IC3, IC5). Note the noise components had spatial distributions associated with non-gray-matter regions (CSF, draining veins). Note also the networks are labeled and ordered as explained in the text. Bottom bar denotes the different stages of the behavioral task. The amplitude of the component time courses reflects the strength of engagement of each independent component over time, in z-scored units. Modulation of these timecourses by emotions is investigated further below.

Further investigation of the spatial maps using dual regression and statistical modeling revealed insight into the specific regions that were functionally co-activated within each IC network. Specifically, modeling of the functional connectivity data with equation 1 revealed a significant interaction between *IC*_*network* and *Region* (p *<* 0.0001; see Table 2 for all stats), suggesting that the pattern of regional functional connectivity was not the same across the four IC networks. Post-hoc analyses with Bonferroni correction revealed patterns of regional functional connectivity that were unique to each IC network. These patterns are visualized in Figures 4 and 5. Whereas both IC0 (“meaning”) and IC1 (“input”) strongly relied on visual areas and temporal areas (lateral occipital, pericalcarine gyrus), IC1 (“input”) differed from IC0 (“meaning”) in its co-activity in the superior temporal gyrus, amygdala, and medial posterior cingulate regions. The IC2 (“response”) network had strong activity in the primary motor regions as well as the anterior cingulate. Finally, the IC4 (“dmn”) network showed strong activity in typical default mode network regions.

**Table 2.** Overview of the ANOVA table from the statistical modeling of the regional functional connectivity data.

|  | Sum Sq | Mean Sq | NumDF | DenDF | F value | Pr(>F) |
| --- | --- | --- | --- | --- | --- | --- |
| Hemisphere | 0.17 | 0.17 | 1.00 | 21155.06 | 6.15 | 0.0131 |
| Headmotion | 0.03 | 0.03 | 1.00 | 19.03 | 0.98 | 0.3346 |
| Euler | 0.07 | 0.07 | 1.00 | 18.97 | 2.53 | 0.1279 |
| IC_network | 26.37 | 8.79 | 3.00 | 21155.54 | 309.64 | < 0.0001 |
| Emotion_condition | 0.09 | 0.04 | 2.00 | 21155.53 | 1.58 | 0.2060 |
| Region | 66.83 | 1.63 | 41.00 | 21155.13 | 57.43 | < 0.0001 |
| IC:cond | 0.50 | 0.08 | 6.00 | 21155.46 | 2.95 | 0.0070 |
| IC:region | 226.84 | 1.84 | 123.00 | 21155.08 | 64.98 | < 0.0001 |
| cond:region | 1.81 | 0.02 | 82.00 | 21155.07 | 0.78 | 0.9313 |
| IC:cond:region | 12.00 | 0.05 | 246.00 | 21155.05 | 1.72 | < 0.0001 |

**Figure 4.**
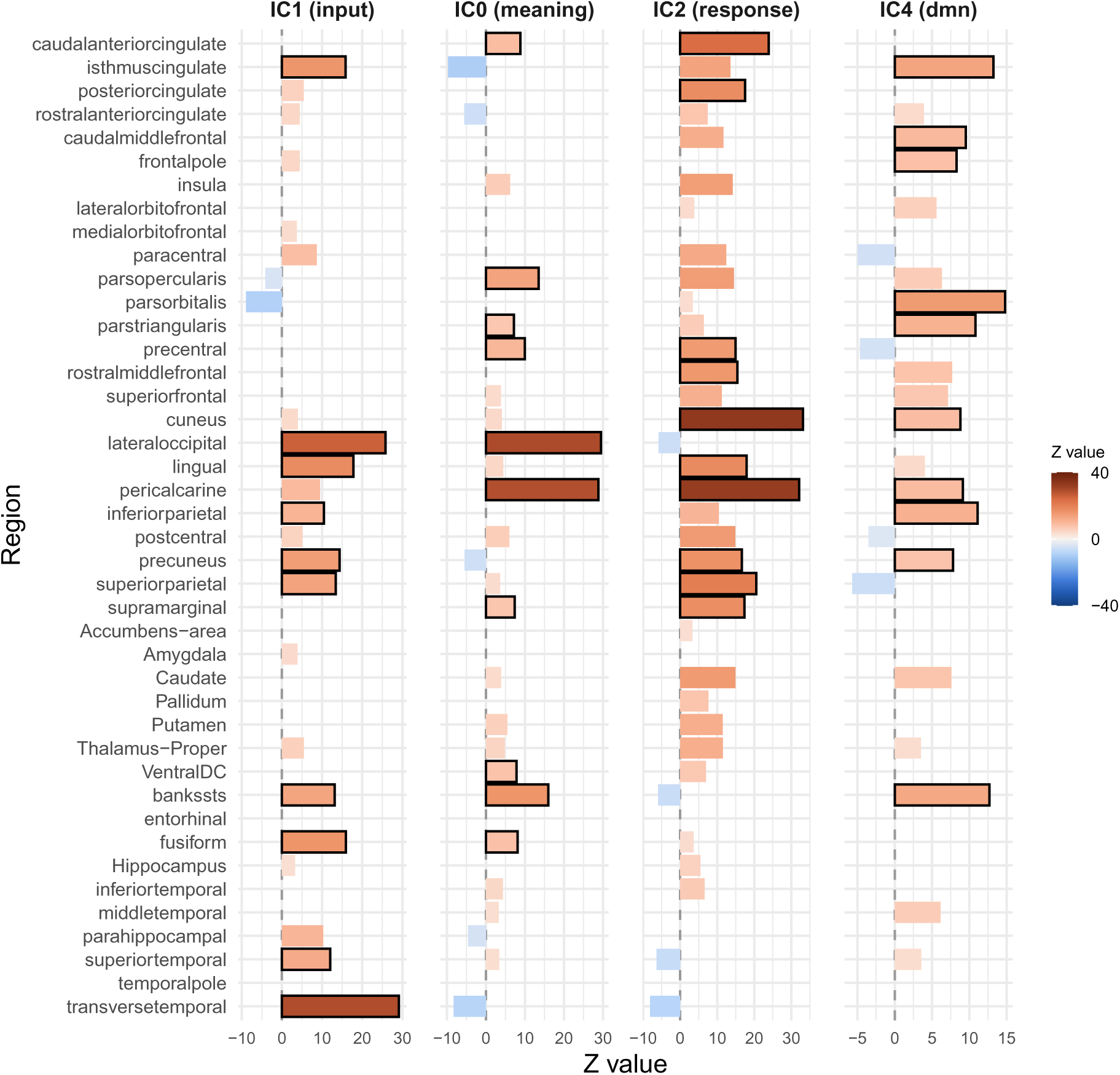
Basic regional functional connectivity in the four large scale networks. Note scale differences between networks. Black bars denote top 10 regions with highest functional connectivity within each network. All results are bonferroni corrected.

**Figure 5.**
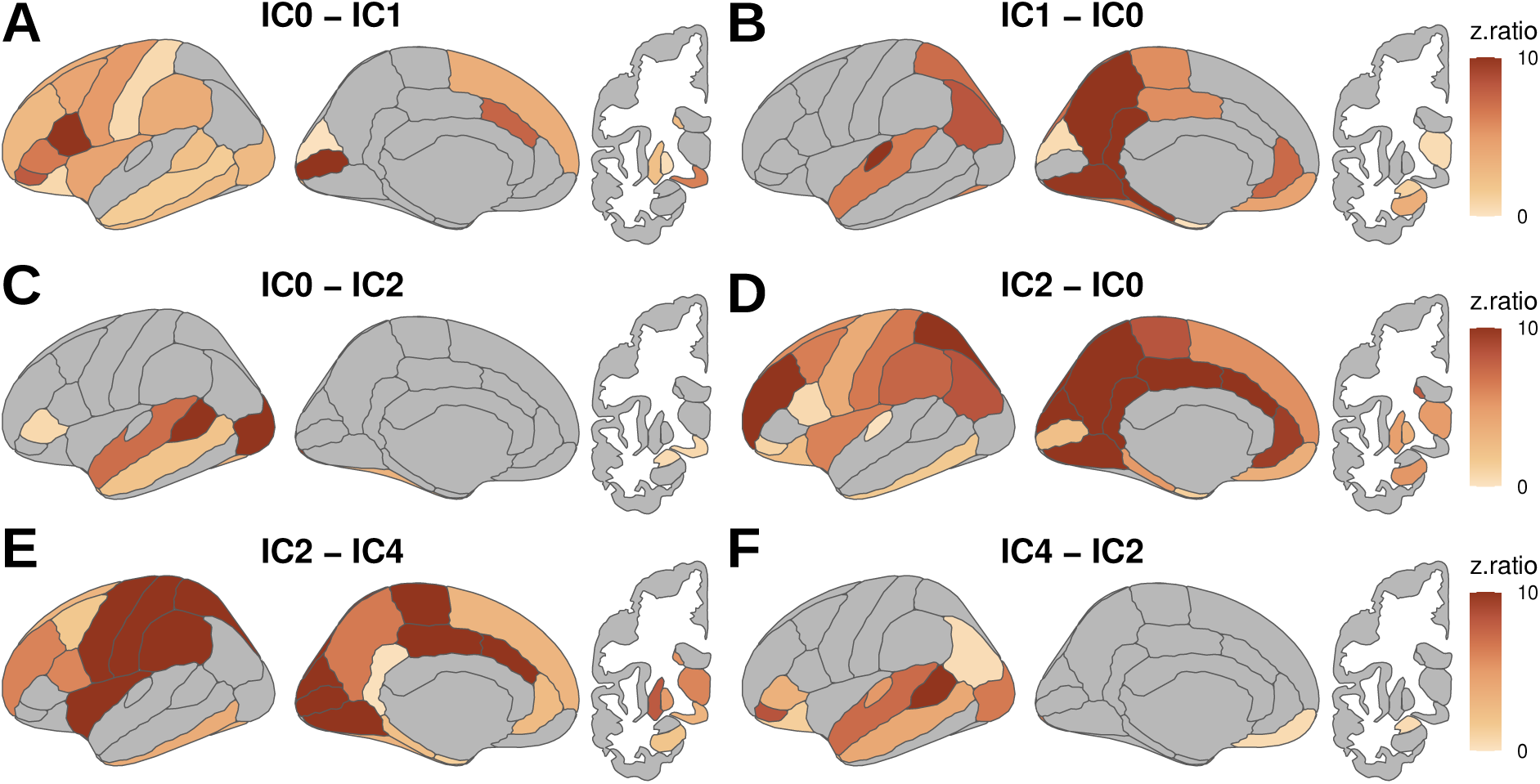
Differences in functional connectivity between IC networks. Note here we focus on three main contrasts to highlight differences between IC0 and IC1, IC0 and IC2 and between IC2 and IC4.

Statistical modeling with equation 1 also revealed a triple interaction between *IC*_*network*, *Region* and *Emotion*_*Condition* (p *<* 0.0001; see Table 2 for all stats), suggesting that the interaction between *IC*_*network* and *Region* outlined above was further modulated by the Emotion Condition. Post-hoc analyses of this three-way interaction revealed different effects of Emotion Condition for different regions within each IC network. As can be seen in Figure 6, discrete emotions modulated regions across the whole brain within all four IC networks. Specifically, discrete emotion modulated functional connectivity in visual, temporal, frontal, parietal and subcortical regions for all four networks.

**Figure 6.**
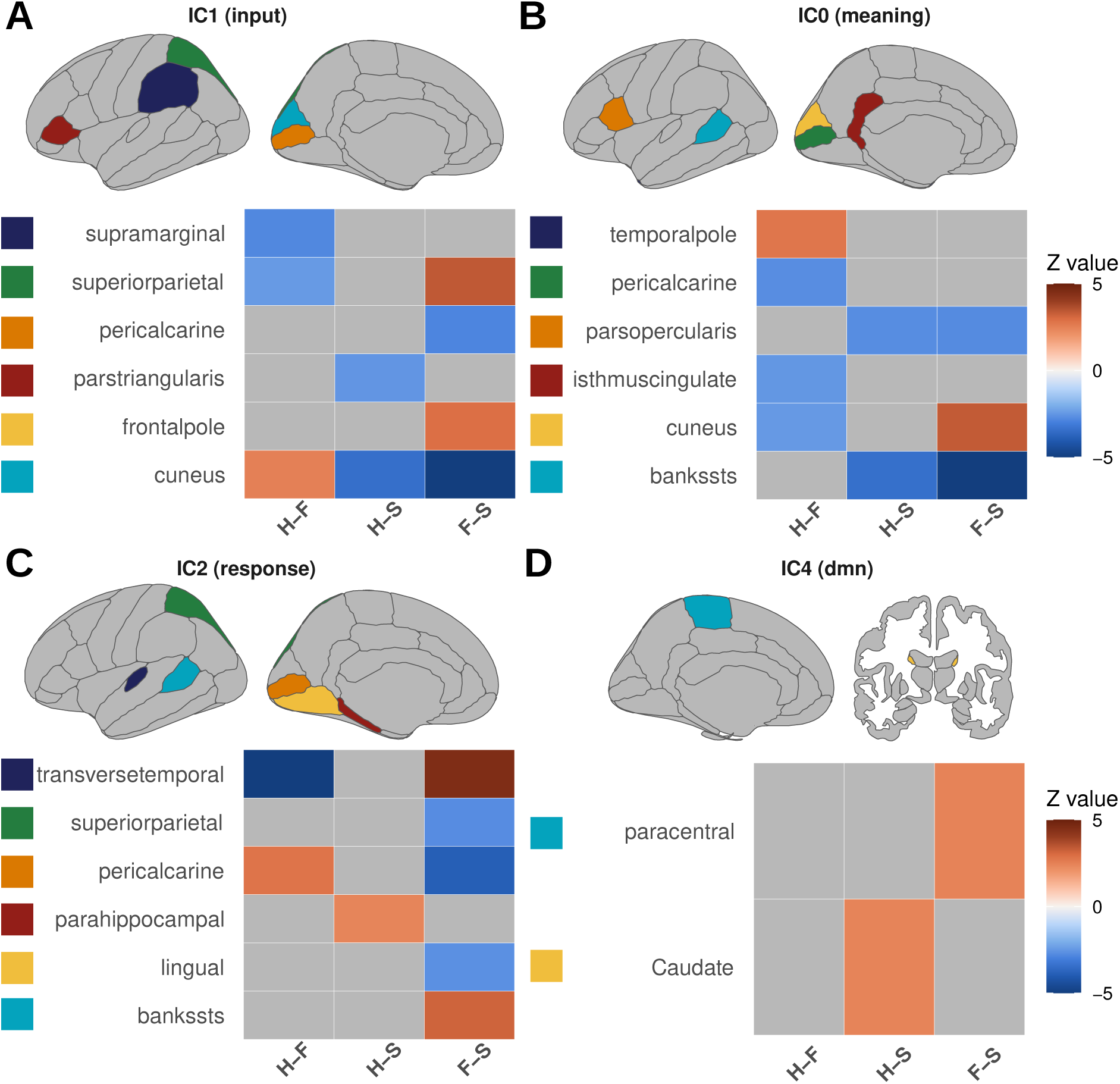
Functional connectivity modulation of emotion conditions across networks. Note all large scale networks involved in the task are modulated by emotions in all major lobes of the brain. H = Happy, F = Fear, S = Sadness. All results are bonferroni corrected.

Curve-fitting of a Gaussian function to the temporal profiles associated with the four IC networks revealed further insight into the temporal properties of the four networks. First, the results revealed relatively high R2 values suggesting that goodness of fit was adequate across the four IC networks (see Figure 7B). In addition, statistical modeling with equation 3 revealed a main effect of *IC*_*network* for all three dependent variables suggesting that peak values, time to peak and durations were not equal for the four IC networks (see Table 3 for details). Post-hoc analyses of *IC*_*network* for each dependent variable revealed that peak values differed between the four networks, where time to peak values were earlier for IC1 (“input”) than for IC0 (“meaning”), and durations were generally shorter for the networks associated in the later aspects of the task (see Figure 7C, D and E for further details).

**Figure 7.**
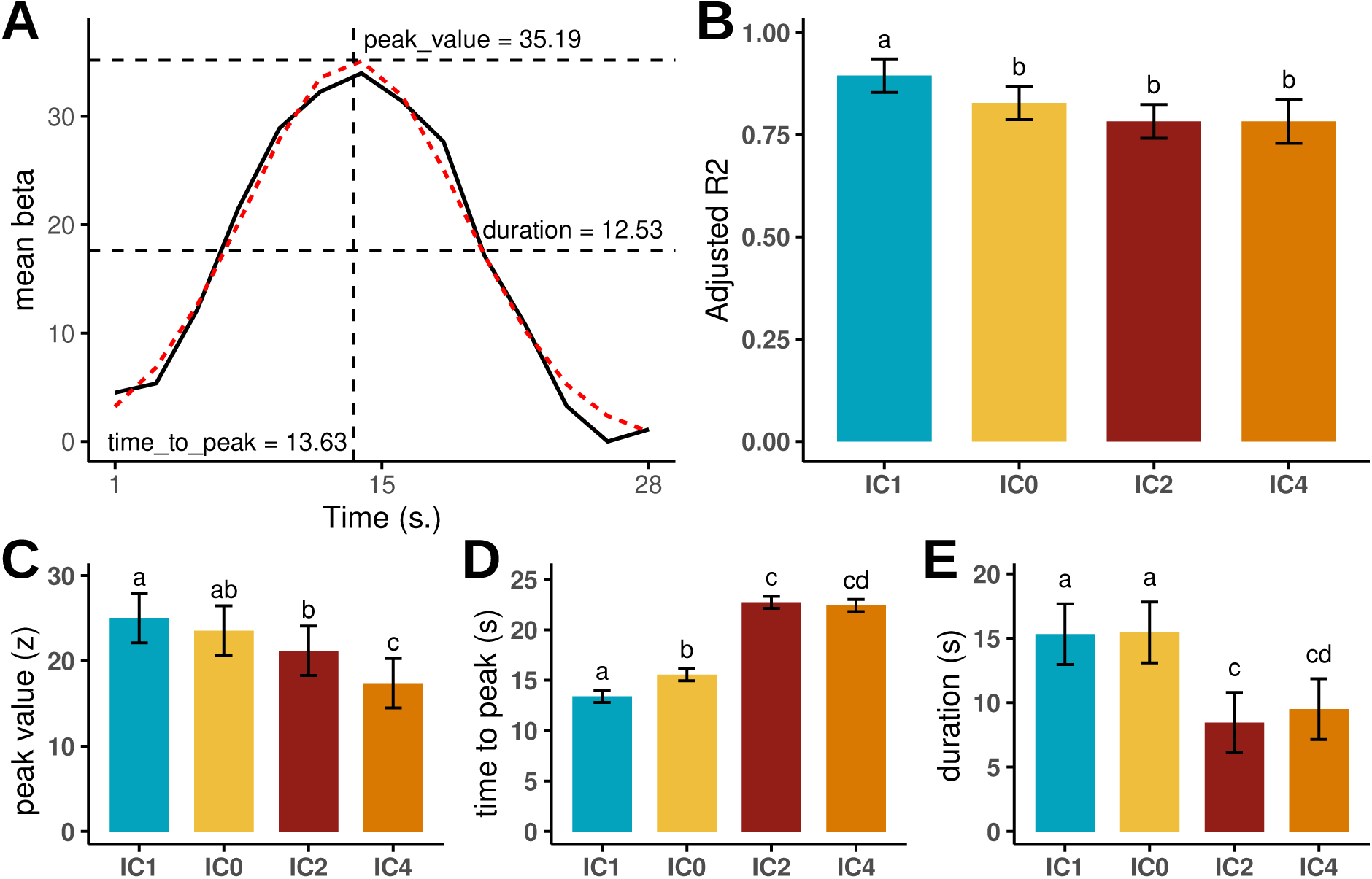
Explanation of curvefitting procedure (A), goodness of fit results from Gaussian curvefitting (B), and graphical overview of how peak value (C), time to peak (D) and duration (E) differ between the four IC networks. Note that goodness of fit values were relatively equal across IC networks, and that there were earlier time to peak values for IC1 compared to IC0. Different letters above bars indicate a significant difference (p < 0.05, bonferroni corrected).

Finally, statistical modeling with equation 3 also revealed an interaction between *IC*_*network* and *Emotion*_*Condition* for all three dependent variables, suggesting that peak value, time to peak and duration values for the happy, sad and fear conditions were not equal across the four IC networks (see Table 3 for details). Post-hoc analyses revealed that the Emotion Condition modulated peak value and duration for IC1 (“input”), peak value and time to peak for IC0 (“meaning”), time to peak for IC2 (“response”) and duration for IC4 (“dmn”; see Figure 8A-C for additional details).

**Table 3.** Overview of the ANOVA table from the statistical modeling of the curvefitting data.

|  | Variable | Sum Sq | Mean Sq | NumDF | DenDF | F value | Pr(>F) |
| --- | --- | --- | --- | --- | --- | --- | --- |
| peak value | RSE | 2583.06 | 2583.06 | 1.00 | 206.16 | 77.37 | < 0.0001 |
|  | IC | 740.74 | 246.91 | 3.00 | 190.48 | 7.40 | 0.0001 |
|  | cond | 1123.70 | 561.85 | 2.00 | 187.37 | 16.83 | < 0.0001 |
|  | IC:cond | 1058.63 | 176.44 | 6.00 | 186.05 | 5.28 | < 0.0001 |
| time to peak | RSE | 1.72 | 1.72 | 1.00 | 205.18 | 1.67 | 0.1981 |
|  | IC | 3403.49 | 1134.50 | 3.00 | 189.38 | 1096.81 | < 0.0001 |
|  | cond | 26.45 | 13.22 | 2.00 | 187.35 | 12.78 | < 0.0001 |
|  | IC:cond | 14.83 | 2.47 | 6.00 | 186.55 | 2.39 | 0.0301 |
| duration | RSE | 9.43 | 9.43 | 1.00 | 160.08 | 3.25 | 0.0734 |
|  | IC | 1443.68 | 481.23 | 3.00 | 195.80 | 165.69 | < 0.0001 |
|  | cond | 24.82 | 12.41 | 2.00 | 191.79 | 4.27 | 0.0153 |
|  | IC:cond | 46.73 | 7.79 | 6.00 | 189.42 | 2.68 | 0.0160 |

**Figure 8.**
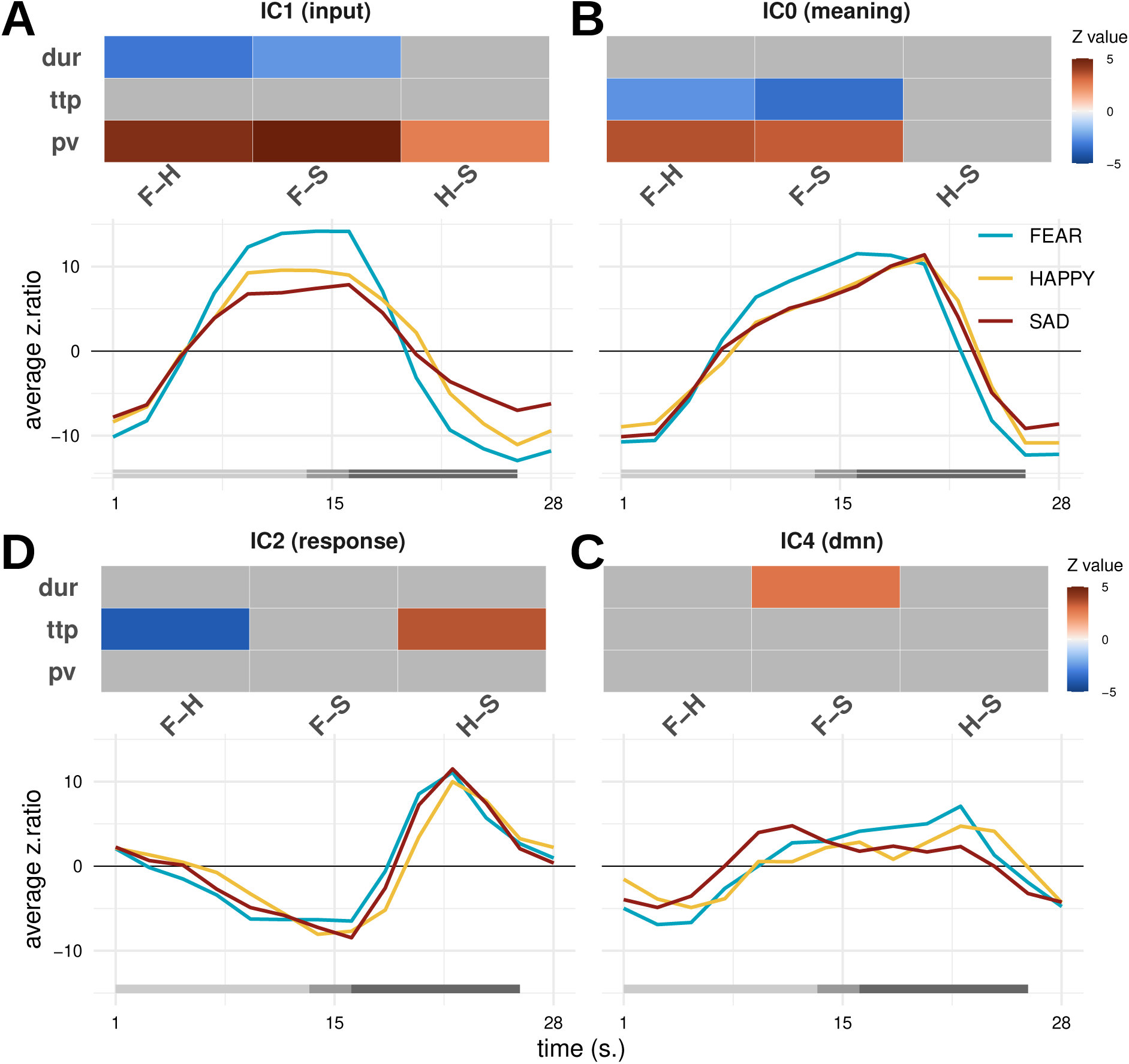
Results of the pairwise contrasts of the Happy (H), Sadness (S) and Fear (F) conditions for the peak value (pv), time to peak (ttp) and duration (dur) across all four IC networks. Only significant (*p* 0.05, bonferroni corrected) values are plotted.

## Discussion

In the current study we relied on time-resolved fMRI to discover the large scale networks associated with a behavioral task that consisted of three stages: watching a movie clip (12.5s), making a decision about the emotion experience by the protagonist in the clip (2.5s) and an intertrial rest period (10s). Group spatial ICA of slice-based fMRI data with increased temporal resolution across the entire trial period (28s) revealed four large scale brain networks that were associated with different aspects of the task. Two networks (IC0 and IC1) were associated with the movie stimulus, while a third network (IC2) activated during the later decision stage, and a fourth network (IC4) was activated throughout the task and showed a peak during the intertrial rest period. Further quantification of the spatial and temporal properties of these networks using dual regression and Gaussian curvefitting showed that the emotional content of the clip modulated the spatial distribution as well as the peak value, time to peak and duration properties of each network.

Brain networks IC0 and IC1 appear to play complementary roles during the movie-watching phase. IC1, which we labeled the “input” network, aligns with aspects of Networks 1 and 2 of Riedel et al. (2018), involving primary visual and auditory processing regions critical for perceiving the movie stimulus (see Figure 4). However, IC1 (“input”) also showed a resemblance to Network 5 of Riedel et al. (2018) with activity in the amygdala, fusiform gyrus and parahippocampal gyrus that was linked to the evaluation of the emotional content of the stimulus. Note that no other network detected in our data consisted of co-activity in these three regions. In addition, time to peak values were earlier in the IC1 (“input”) network than in all other networks (see Figure 7D). The combined spatial and temporal properties of this network therefore suggests that this network is likely responsible for processing the visual and auditory input stream as well as the evaluation of the emotional content of the stimulus.

On the other hand, although also associated with the movie stimulus, the IC0 (“meaning”) network appeared to have spatial and temporal properties that were distinct from the IC1 (“input”) network. Specifically, IC0 (“meaning”) had increased co-activity in lateral frontal and middle temporal areas that are typically associated with semantic processing (see Figure 5; Huth et al., 2016; Visser et al., 2012). In addition, IC0 (“meaning”) aligned with Network 3 of Riedel et al. (2018) and appeared to contain regions associated with the salience network such as the insula and the anterior cingulate gyrus (Seeley, 2019), suggesting an attentional component. Importantly, even though the IC0 (“meaning”) network activated in response to the movie stimulus, its time to peak values lagged about 2s behind those of IC1 (“input”; see Figure 7). As was pointed out in the Introduction, the movie clips showed scenes whose intended meaning could not be discerned immediately, but instead required additional processing that lagged behind the input stream. As before, the combined spatial and temporal characteristics of this network allow us to interpret this network as involved in the processing of the meaning associated with the input stream.

In addition, a third network (IC2, “response”) showed temporal activity with a peak around 20s and can therefore be associated with the response stage (see Figure 3). The spatial properties of this network showed activity in cortical and subcortical regions associated with processing the visual input (the discrete emotion words), decision making associated regions like the caudate nucleus (Grahn, Parkinson, & Owen, 2008), motor generation regions like the precentral gyrus, and task general attention related regions like the insula and the anterior cingulate cortex (Seeley, 2019). This therefore suggests that this network represents the combined emotion decision and motor output network. Note that with increased temporal resolution (and potentially restricting behavioral variance in response latency) one might be able to separate the decision and the motor output components of this network. Finally, the fourth network (IC4, “dmn”) showed an alignment with Network 4 of Riedel et al. (2018) and had spatial properties consistent with those of the default mode network (Buckner, Andrews-Hanna, & Schacter, 2008). Although the exact role of the default mode network in emotion processing is currently debated (Satpute & Lindquist, 2019), it is noteworthy that the network showed some co-activity throughout the task and had a peak during the rest period (see Figure 3), and showed activity in lateral prefrontal regions (see Figure 4). These spatial and temporal properties are consistent with an interpretation in terms of the regulation, mind-wandering and/or self-referential processing of the emotions (Gross, 2015).

Both the spatial and temporal properties of the four identified networks were influenced by the emotional content of the movie clips. Specifically, in IC1 (“input”), regions in the visual, parietal, and frontal lobes showed differential activation patterns across happy, fear, and sad clips. Temporal dynamics, including the duration and peak values of the responses, also varied between these emotional conditions (Figure 6A and 8A). Similar modulations were observed for IC0 (“meaning”), IC2 (“response”), and IC4 (“dmn”), where emotional content influenced activations in visual, temporal, and frontal regions, as well as temporal metrics such as response duration, time to peak, and peak amplitude. Notably, the time-to-peak differences in IC2 (“response”) align closely with response latency results, with sad trials showing faster response latencies and earlier peak times (compare Figures 1D and 8D). Critically, the same ordering between response latencies and the fMRI time-to-peak metric provides convergent temporal validity for using BOLD dynamics as a window into the speed of affective computations. Nevertheless, these results should not be taken as evidence that processing sad emotions is generally faster than other emotions; Other studies have demonstrated fast processing of fearful stimuli (Bo et al., 2022; Grootswagers, Kennedy, Most, & Carlson, 2020; Méndez-Bértolo et al., 2016), and specific aspects of the current experimental paradigm likely influenced the speed of emotional processing. In general, the current findings suggest that emotional stimuli elicit widespread effects on brain activity, impacting not only the processing of emotion-laden input stimuli but also the spatial and temporal dynamics of broader neural responses.

Taken together, the results of this study paint a comprehensive picture of how the brain orchestrates spatially and temporally distinct networks to process emotional stimuli and execute task-relevant behaviors. The findings highlight a dynamic interplay between sensory processing, semantic interpretation, decision-making, and emotional regulation, all of which unfold within a distributed but highly coordinated network architecture. Although the networks demonstrated separable time courses, their temporal dynamics were largely overlapping, suggesting that processes such as those mediated by IC1 (“input”) and IC0 (“meaning”) likely operate in parallel rather than in isolation. This parallel engagement implies that cross-talk between networks is a critical feature of task-related brain activity (Cole et al., 2013), particularly during complex cognitive and emotional tasks. For example, the interplay between sensory-driven processing in IC1 and the semantic and attentional mechanisms in IC0 may be necessary for the integrated perception of emotionally salient stimuli. How such cross-talk is mediated remains an open question, representing an intriguing avenue for future research. In the decision phase, IC2 (“response”) reveals how emotion processing transitions from perception to cognitive evaluation, involving a convergence of sensory, attentional, and motor networks to guide task execution. Finally, IC4, (“dmn”) associated with the default mode network, demonstrates the presence of reflective, mind-wandering and/or regulatory processes (Gross, 2015), particularly during rest but with influence throughout the task (see Figure 3). These findings underscore the dynamic nature of brain activity, where emotional stimuli engage overlapping networks that continuously integrate sensory input, semantic processing, decision-making, and regulation. This fluid coordination reveals how emotions are adaptively represented and processed across time and space (John et al., 2022; Pessoa & McMenamin, 2017; Waugh et al., 2015).

Our study has several limitations that warrant discussion. First, the interpretation of BOLD signal dynamics is inherently influenced by the vascular properties of the underlying tissue (Ogawa et al., 1992). This limitation has been discussed primarily in the context of studies that have attempted to determine the temporal sequence of activation of different brain regions where such differences could be confounded by variations in vascular responsiveness (Aguirre, Zarahn, & D’Esposito, 1998; Kim, Richter, & Ugurbil, 1997; Menon, Luknowsky, & Gati, 1998). However, it is important to note that the temporal properties of the signal we analyzed do not represent those of individual brain regions but the aggregate activity of many regions working in concert, reducing the likelihood of significant confounding effects. A further limitation is that we did not require participants to directly regulate their emotions which complicates an interpretation in terms of emotion regulation (Gross, 2015). Likewise, the current research design only used three main emotional categories. Whether our conclusions hold for other emotions and emotion types remains to be studied in future studies. Another general limitation of the results presented here concerns the influence of the number of dimensions chosen in group spatial ICA on the spatial distribution of the observed large-scale networks. It is well-documented that increasing the number of dimensions can lead to the “splitting” of networks into finer components, while fewer dimensions may overgeneralize distinct functional patterns (Beckmann, 2012). This presents a challenge, as there is no a priori standard for the “correct” spatial distribution of task-related networks, especially for complex tasks like watching movie clips (Di, Gohel, Kim, & Biswal, 2013; Dosenbach, Raichle, & Gordon, 2025; Huang, De Brigard, Cabeza, & Davis, 2024). In this sense, task-based fMRI studies of large-scale networks are currently in a similar position to early resting-state fMRI research, where the precise spatial distributions of networks were still being defined (e.g., Yeo et al., 2011). To address this, a concerted effort is needed to map the large-scale networks underlying diverse tasks, enabling more informed choices about dimensionality and mitigating the uncertainty in network characterization.

Finally, it is worth emphasizing that our approach bears a relationship to what is often termed “dynamic functional connectivity” (Gonzalez-Castillo & Bandettini, 2018; Hutchison et al., 2013). In particular, our results demonstrate how whole-brain functional connectivity maps shift over time in response to different components of a complex, emotion-based behavioral task (Figure 3). Although both event-related fMRI and group spatial ICA have been employed extensively in prior studies, combining them as in the current study is less common (Janssen & Mendieta, 2020; Long et al., 2013). The particular slice-based event-related approach has been shown to offer enhanced temporal precision and greater flexibility in choosing different temporal resolutions than more traditional signal extraction schemes (Janssen et al., 2018). Indeed, when we analyzed our data at lower temporal resolution (i.e., at the 3 s TR level), some components with insufficient temporal separation (such as IC1 and IC0) became conflated, highlighting the importance of higher-resolution signal extraction in distinguishing these networks (Calhoun, Adali, Pearlson, & Pekar, 2001). Moreover, we used standard group spatial ICA methods, which have been widely applied to resting-state data, to derive task-based whole-brain functional connectivity maps. This unified approach for finding networks in task-based and resting-state data may facilitate comparisons between networks obtained from different modalities (Gonzalez-Castillo & Bandettini, 2018). Taken together, these findings demonstrate the value of combining slice-based event-related fMRI with group spatial ICA to characterize how functional connectivity patterns evolve over time. Future research may further explore the interplay of the temporal resolution of the extracted signal, group spatial ICA, and alternative approaches, such as FIR analysis (Josephs et al., 1997; Windischberger et al., 2008), to illuminate the spatiotemporal characteristics of dynamic network changes during complex tasks.

To conclude, the present study addressed three key questions: whether distinct whole-brain networks underlie the different stages of our emotional movie watching and decision task, whether their spatial distributions vary with emotional content, and whether their temporal dynamics—including time-to-peak and duration—are sensitive to emotion. Our results revealed four large-scale networks whose activity patterns not only differentiated task stages but revealed a separation between a network related to processing affective input and understanding the meaning of this input. Importantly, the spatial and temporal features of all four networks varied as a function of the emotional content of the stimuli. These results move beyond static, region-based models and suggest emotion reflects dynamic, distributed interactions among multiple networks (K. A. Lindquist et al., 2012; Pessoa, 2017). In turn, our findings underscore the utility of examining temporal metrics to capture subtle nuances of emotional processing that may remain undetectable using standard static analyses. These temporal properties may be especially relevant for clinical studies in which certain emotional disorders arise due to disturbances in the temporal regulation of emotion (e.g., dwelling too long on certain thoughts; Kuppens & Verduyn, 2017; Waugh et al., 2015). The current study’s approach for quantifying these temporal properties can therefore offer valuable insights, both for basic science and for the development of novel clinical tools.

## Acknowledgments

The authors would like to thank the Servicio de Resonancia Magnética para Investigaciones Biomédicas (SRMIB - SEGAI) at the University of La Laguna for their help.

## Notes

### Competing Interest Statement

The authors have declared no competing interest.

### Summary of Updates

This version of the manuscript was revised in accordance with the journal reviewer requests.

### Funding

- Ministerio de Ciencia, Innovación y Universidades: PID2021-127611NB-I00

## References

Adolphs, R. (2002). Recognizing emotion from facial expressions: psychological and neurological mechanisms. Behavioral and cognitive neuroscience reviews, 1 (1), 21–62.

Aguirre, G. K., Zarahn, E., & D’Esposito, M. (1998). The variability of human, bold hemodynamic responses. Neuroimage, 8 (4), 360–369.

Bates, D., Mächler, M., Bolker, B., & Walker, S. (2015). Fitting linear mixed-effects models using lme4. Journal of Statistical Software, 67 (1), 1–48. doi: 10.18637/jss.v067.i01

Beckmann, C. F. (2012). Modelling with independent components. Neuroimage, 62 (2), 891–901.

Beckmann, C. F., Mackay, C. E., Filippini, N., Smith, S. M., et al. (2009). Group comparison of resting-state fmri data using multi-subject ica and dual regression. Neuroimage, 47 (Suppl 1), S148.

Beckmann, C. F., & Smith, S. M. (2004). Probabilistic independent component analysis for functional magnetic resonance imaging. IEEE transactions on medical imaging, 23 (2), 137–152.

Bo, K., Cui, L., Yin, S., Hu, Z., Hong, X., Kim, S., … Ding, M. (2022). Decoding the temporal dynamics of affective scene processing. NeuroImage, 261, 119532.

Buckner, R. L., Andrews-Hanna, J. R., & Schacter, D. L. (2008). The brain’s default network: anatomy, function, and relevance to disease. Annals of the new York Academy of Sciences, 1124 (1), 1–38.

Calhoun, V. D., Adali, T., Pearlson, G., & Pekar, J. J. (2001). Spatial and temporal independent component analysis of functional mri data containing a pair of task-related waveforms. Human brain mapping, 13 (1), 43–53.

Cole, M. W., Reynolds, J. R., Power, J. D., Repovs, G., Anticevic, A., & Braver, T. S. (2013). Multi-task connectivity reveals flexible hubs for adaptive task control. Nature neuroscience, 16 (9), 1348–1355.

Desikan, R. S., Ségonne, F., Fischl, B., Quinn, B. T., Dickerson, B. C., Blacker, D., … others (2006). An automated labeling system for subdividing the human cerebral cortex on mri scans into gyral based regions of interest. Neuroimage, 31 (3), 968–980.

Di, X., Gohel, S., Kim, E. H., & Biswal, B. B. (2013). Task vs. rest—different network configurations between the coactivation and the resting-state brain networks. Frontiers in human neuroscience, 7, 493.

Dosenbach, N. U., Raichle, M. E., & Gordon, E. M. (2025). The brain’s action-mode network. Nature Reviews Neuroscience, 1–11.

Duann, J.-R., Jung, T.-P., Kuo, W.-J., Yeh, T.-C., Makeig, S., Hsieh, J.-C., & Sejnowski, T. J. (2002, apr). Single-trial variability in event-related BOLD signals. NeuroImage, 15 (4), 823–835. Retrieved from 10.1006/nimg.2001.1049 doi: 10.1006/nimg.2001.1049

Elzhov, T. V., Mullen, K. M., Spiess, A.-N., & Bolker, B. (2022). minpack.lm: R interface to the levenberg-marquardt nonlinear least-squares algorithm [Computer software manual]. Retrieved from https://CRAN.R-project.org/package=minpack.lm (R package version X.X)

Etkin, A., Büchel, C., & Gross, J. J. (2015). The neural bases of emotion regulation. Nature reviews neuroscience, 16 (11), 693–700.

Finn, E. S., & Bandettini, P. A. (2021). Movie-watching outperforms rest for functional connectivity-based prediction of behavior. NeuroImage, 235, 117963.

Fischl, B. (2012). Freesurfer. Neuroimage, 62 (2), 774–781.

Glasser, M. F., Sotiropoulos, S. N., Wilson, J. A., Coalson, T. S., Fischl, B., Andersson, J. L., … others (2013). The minimal preprocessing pipelines for the human connectome project. Neuroimage, 80, 105–124.

Gonzalez-Castillo, J., & Bandettini, P. A. (2018). Task-based dynamic functional connectivity: Recent findings and open questions. Neuroimage, 180, 526–533.

Grahn, J. A., Parkinson, J. A., & Owen, A. M. (2008). The cognitive functions of the caudate nucleus. Progress in neurobiology, 86 (3), 141–155.

Griffanti, L., Salimi-Khorshidi, G., Beckmann, C. F., Auerbach, E. J., Douaud, G., Sexton, C. E., … others (2014). Ica-based artefact removal and accelerated fmri acquisition for improved resting state network imaging. Neuroimage, 95, 232–247.

Grootswagers, T., Kennedy, B. L., Most, S. B., & Carlson, T. A. (2020). Neural signatures of dynamic emotion constructs in the human brain. Neuropsychologia, 145, 106535.

Gross, J. J. (2015). The extended process model of emotion regulation: Elaborations, applications, and future directions. Psychological inquiry, 26 (1), 130–137.

Huang, S., De Brigard, F., Cabeza, R., & Davis, S. W. (2024). Connectivity analyses for task-based fmri. Physics of Life Reviews.

Hutchison, R. M., Womelsdorf, T., Allen, E. A., Bandettini, P. A., Calhoun, V. D., Corbetta, M., … others (2013). Dynamic functional connectivity: promise, issues, and interpretations. Neuroimage, 80, 360–378.

Huth, A. G., De Heer, W. A., Griffiths, T. L., Theunissen, F. E., & Gallant, J. L. (2016). Natural speech reveals the semantic maps that tile human cerebral cortex. Nature, 532 (7600), 453–458.

Jääskeläinen, I. P., Sams, M., Glerean, E., & Ahveninen, J. (2021). Movies and narratives as naturalistic stimuli in neuroimaging. NeuroImage, 224, 117445.

Janssen, N., Hernández-Cabrera, J. A., & Foronda, L. E. (2018). Improving the signal detection accuracy of functional magnetic resonance imaging. NeuroImage, 176, 92–109.

Janssen, N., & Mendieta, C. C. R. (2020). The dynamics of speech motor control revealed with time-resolved fmri. Cerebral Cortex, 30 (1), 241–255.

Jenkinson, M., Bannister, P., Brady, M., & Smith, S. (2002). Improved optimization for the robust and accurate linear registration and motion correction of brain images. Neuroimage, 17 (2), 825–841.

John, Y. J., Sawyer, K. S., Srinivasan, K., Müller, E. J., Munn, B. R., & Shine, J. M. (2022). It’s about time: Linking dynamical systems with human neuroimaging to understand the brain. Network Neuroscience, 6 (4), 960–979.

Josephs, O., Turner, R., & Friston, K. (1997). Event-related fmri. Human brain mapping, 5 (4), 243–248.

Kim, S.-G., Richter, W., & Ugurbil, K. (1997). Limitations of temporal resolution in functional mri. Magnetic resonance in medicine, 37 (4), 631–636.

Kober, H., Barrett, L. F., Joseph, J., Bliss-Moreau, E., Lindquist, K., & Wager, T. D. (2008). Functional grouping and cortical–subcortical interactions in emotion: a meta-analysis of neuroimaging studies. Neuroimage, 42 (2), 998–1031.

Kruggel, F., & von Cramon, D. Y. (1999). Modeling the hemodynamic response in single-trial functional mri experiments. Magnetic Resonance in Medicine, 42 (4), 787–797.

Kruggel, F., Zysset, S., & von Cramon, D. Y. (2000). Nonlinear regression of functional mri data: an item recognition task study. NeuroImage, 12 (2), 173–183.

Kuppens, P., & Verduyn, P. (2017). Emotion dynamics. Current Opinion in Psychology, 17, 22–26.

Kuznetsova, A., Brockhoff, P. B., Christensen, R. H., et al. (2017). lmertest package: tests in linear mixed effects models. Journal of statistical software, 82 (13), 1–26.

LaBar, K. S., LeDoux, J. E., Spencer, D. D., & Phelps, E. A. (1995). Impaired fear conditioning following unilateral temporal lobectomy in humans. Journal of neuroscience, 15 (10), 6846–6855.

Lenth, R., Singmann, H., Love, J., Buerkner, P., & Herve, M. (2018). Emmeans: Estimated marginal means, aka least-squares means. R package version, 1 (1), 3.

Lindquist, K. A., Wager, T. D., Kober, H., Bliss-Moreau, E., & Barrett, L. F. (2012). The brain basis of emotion: a meta-analytic review. Behavioral and brain sciences, 35 (3), 121–143.

Lindquist, M. A., Loh, J. M., Atlas, L. Y., & Wager, T. D. (2009). Modeling the hemodynamic response function in fmri: efficiency, bias and mis-modeling. Neuroimage, 45 (1), S187–S198.

Liu, M., Liu, C. H., Zheng, S., Zhao, K., & Fu, X. (2021). Reexamining the neural network involved in perception of facial expression: A meta-analysis. Neuroscience & Biobehavioral Reviews, 131, 179–191.

Long, Z., Li, R., Wen, X., Jin, Z., Chen, K., & Yao, L. (2013). Separating 4d multi-task fmri data of multiple subjects by independent component analysis with projection. Magnetic Resonance Imaging, 31 (1), 60–74.

Meer, J. N. v. d., Breakspear, M., Chang, L. J., Sonkusare, S., & Cocchi, L. (2020). Movie viewing elicits rich and reliable brain state dynamics. Nature communications, 11 (1), 5004.

Méndez-Bértolo, C., Moratti, S., Toledano, R., Lopez-Sosa, F., Martínez-Alvarez, R., Mah, Y. H., … Strange, B. A. (2016). A fast pathway for fear in human amygdala. Nature neuroscience, 19 (8), 1041–1049.

Menon, R. S., Luknowsky, D. C., & Gati, J. S. (1998). Mental chronometry using latency-resolved functional mri. Proceedings of the National Academy of Sciences, 95 (18), 10902–10907.

Minka, T. (2000). Automatic choice of dimensionality for pca. Advances in neural information processing systems, 13 .

Morgenroth, E., Vilaclara, L., Muszynski, M., Gaviria, J., Vuilleumier, P., & Van De Ville, D. (2023). Probing neurodynamics of experienced emotions—a hitchhiker’s guide to film fmri. Social Cognitive and Affective Neuroscience, 18 (1), nsad063.

Murphy, F. C., Nimmo-Smith, I., & Lawrence, A. D. (2003). Functional neuroanatomy of emotions: a meta-analysis. Cognitive, affective, & behavioral neuroscience, 3 (3), 207–233.

Ochsner, K. N., Bunge, S. A., Gross, J. J., & Gabrieli, J. D. (2002). Rethinking feelings: an fmri study of the cognitive regulation of emotion. Journal of cognitive neuroscience, 14 (8), 1215–1229.

Ochsner, K. N., & Gross, J. J. (2008). Cognitive emotion regulation: Insights from social cognitive and affective neuroscience. Current directions in psychological science, 17 (2), 153–158.

Ogawa, S., Tank, D. W., Menon, R., Ellermann, J. M., Kim, S. G., Merkle, H., & Ugurbil, K. (1992). Intrinsic signal changes accompanying sensory stimulation: functional brain mapping with magnetic resonance imaging. Proceedings of the National Academy of Sciences, 89 (13), 5951–5955.

Olman, C. A., Davachi, L., & Inati, S. (2009). Distortion and signal loss in medial temporal lobe. PloS one, 4 (12), e8160.

Palomero-Gallagher, N., & Amunts, K. (2022). A short review on emotion processing: a lateralized network of neuronal networks. Brain Structure and Function, 227 (2), 673–684.

Parker, D., Liu, X., & Razlighi, Q. R. (2017). Optimal slice timing correction and its interaction with fmri parameters and artifacts. Medical image analysis, 35, 434–445.

Pessoa, L. (2017). A network model of the emotional brain. Trends in cognitive sciences, 21 (5), 357–371.

Pessoa, L., & McMenamin, B. (2017). Dynamic networks in the emotional brain. The Neuroscientist, 23 (4), 383–396.

Phan, K. L., Wager, T., Taylor, S. F., & Liberzon, I. (2002). Functional neuroanatomy of emotion: a meta-analysis of emotion activation studies in pet and fmri. Neuroimage, 16 (2), 331–348.

Riedel, M. C., Yanes, J. A., Ray, K. L., Eickhoff, S. B., Fox, P. T., Sutherland, M. T., & Laird, A. R. (2018). Dissociable meta-analytic brain networks contribute to coordinated emotional processing. Human brain mapping, 39 (6), 2514–2531.

Rosen, A. F., Roalf, D. R., Ruparel, K., Blake, J., Seelaus, K., Villa, L. P., … others (2018). Quantitative assessment of structural image quality. Neuroimage, 169, 407–418.

Saarimäki, H. (2021). Naturalistic stimuli in affective neuroimaging: A review. Frontiers in human neuroscience, 15, 675068.

Satpute, A. B., & Lindquist, K. A. (2019). The default mode network’s role in discrete emotion. Trends in cognitive sciences, 23 (10), 851–864.

Seeley, W. W. (2019). The salience network: a neural system for perceiving and responding to homeostatic demands. Journal of Neuroscience, 39 (50), 9878–9882.

Sladky, R., Friston, K. J., Tröstl, J., Cunnington, R., Moser, E., & Windischberger, C. (2011). Slice-timing effects and their correction in functional mri. Neuroimage, 58 (2), 588–594.

Smith, S. M., Fox, P. T., Miller, K. L., Glahn, D. C., Fox, P. M., Mackay, C. E., … others (2009). Correspondence of the brain’s functional architecture during activation and rest. Proceedings of the national academy of sciences, 106 (31), 13040–13045.

Sonkusare, S., Breakspear, M., & Guo, C. (2019). Naturalistic stimuli in neuroscience: critically acclaimed. Trends in cognitive sciences, 23 (8), 699–714.

Vemuri, K., & Surampudi, B. R. (2015). Evidence of stimulus correlated empathy modes–group ica of fmri data. Brain and cognition, 94, 32–43.

Visser, M., Jefferies, E., Embleton, K. V., & Lambon Ralph, M. A. (2012). Both the middle temporal gyrus and the ventral anterior temporal area are crucial for multimodal semantic processing: distortion-corrected fmri evidence for a double gradient of information convergence in the temporal lobes. Journal of cognitive neuroscience, 24 (8), 1766–1778.

Wallenwein, L. A., Schmidt, S. N., Hass, J., & Mier, D. (2024). Cross-modal decoding of emotional expressions in fmri—cross-session and cross-sample replication. Imaging Neuroscience, 2, 1–15.

Waugh, C. E., Shing, E. Z., & Avery, B. M. (2015). Temporal dynamics of emotional processing in the brain. Emotion Review, 7 (4), 323–329.

Westermann, R., Spies, K., Stahl, G., & Hesse, F. W. (1996). Relative effectiveness and validity of mood induction procedures: A meta-analysis. European Journal of social psychology, 26 (4), 557–580.

Windischberger, C., Cunnington, R., Lamm, C., Lanzenberger, R., Langenberger, H., Deecke, L., … Moser, E. (2008). Time-resolved analysis of fmri signal changes using brain activation movies. Journal of neuroscience methods, 169 (1), 222–230.

Xu, S., Zhang, Z., Li, L., Zhou, Y., Lin, D., Zhang, M., … others (2023). Functional connectivity profiles of the default mode and visual networks reflect temporal accumulative effects of sustained naturalistic emotional experience. NeuroImage, 269, 119941.

Yeo, B. T., Krienen, F. M., Sepulcre, J., Sabuncu, M. R., Lashkari, D., Hollinshead, M., … others (2011). The organization of the human cerebral cortex estimated by intrinsic functional connectivity. Journal of neurophysiology.

Zemeckis, R., Hanks, T., Wright, R., Sinise, G., Williamson, M., Field, S., & Groom, W. (1994). Forrest gump. Paramount Pictures Los Angeles.

Zhou, F., Zhao, W., Qi, Z., Geng, Y., Yao, S., Kendrick, K. M., … Becker, B. (2021). A distributed fmri-based signature for the subjective experience of fear. Nature communications, 12 (1), 6643.

